# A Four Electrode Method to Study Dynamics of Ion Activity and Transport in Skeletal Muscle Fibers

**DOI:** 10.1101/635490

**Authors:** Judith A. Heiny, Stephen C. Cannon, Marino DiFranco

## Abstract

Ion movements across biological membranes, driven by electrochemical gradients or active transport mechanisms, control essential cell functions. Membrane ion movements can manifest as electrogenic currents or electroneutral fluxes, and either process can alter the extracellular and/or intracellular concentration of the transported ion(s). Classical electrophysiological methods allow accurate measurement of membrane ion movements when the transport mechanism produces a net ionic current; however, they cannot directly measure electroneutral fluxes and do not detect any accompanying change in intracellular ion concentrations.

Here, we developed a method for simultaneously measuring ion movements and the accompanying dynamic changes in intracellular ion concentration(s) in intact skeletal muscle fibers under voltage– or current clamp in real time. The method combines a two-microelectrode voltage-clamp with ion-selective and reference microelectrodes (4 electrode system). We validate the electrical stability of the system and the viability of the preparation for periods of approximately 1 h. We demonstrate the power of this method with measurements of intracellular Cl^-^, H^+^, and Na^+^ to show: 1) voltage-dependent redistribution of Cl^-^ ions; 2) intracellular pH changes induced by changes in extracellular pCO_2_; and 3) electroneutral and electrogenic Na^+^ movements controlled by the Na,K-ATPase. The method is useful for studying a range of transport mechanisms in many cell types, particularly when the transmembrane ion movements are electrically silent and/or when the transport activity measurably changes the intracellular activity of a transported ion.

## INTRODUCTION

Ion movements across biological membranes control essential cell functions. A wide range of electrophysiological approaches exist to measure ion movements by membrane channels and pumps with good sensitivity and time resolution. However, not all biologically-important membrane transport processes can be studied using these approaches. Existing electrophysiological techniques detect transmembrane ion movements only when the transport mechanism produces a net change in membrane potential (e.g. transport by ion channels or electrogenic pumps) or a net charge transfer across the membrane. Consequently, they do not detect electroneutral transport processes (carriers, exchangers, et al.). In addition, it is often the change in intracellular concentration of a transported ion that determines the cellular response. However, this dynamic information is lost when the intracellular concentration is measured under steady-state conditions at an end point of the measurement, as is commonly done.

Here, we describe a method to simultaneously measure transmembrane ion movements and the accompanying transient changes in intracellular ion concentration (or, more importantly, changes in ion activity). The activity of an ion, rather than the total (bound + free) concentration, is important because it determines the membrane potential, equilibrium potentials, and other parameters that determine the cellular response (Tsien 1989).

Ion-selective micro-electrodes (ISM), in particular liquid membrane ISMs, represent a highly-sensitive method for measuring the intracellular activity of physiologically-relevant ions (Kessler, Clark et al. 1976, Armstrong and Garcia-Diaz 1980, Mooney, Lyall et al. 1988). An ISM is constructed by filling the tip of a silanized glass micropipette with a water-immiscible organic phase doped with a specific lipophilic ionophore, and filling the rest of the micropipette with saline containing the ion of interest. When in contact with an aqueous media (e.g. a solution, the cytoplasm) containing the ion of interest, a voltage is generated across the organic phase, *V*_*ion*_, which typically shows a Nernstian dependence on ion activity. When impaled in a cell, the readout of an ISM (*V*_*ISM*_) equals the arithmetic sum of *V*_*ion*_ and the membrane potential (*V*_*m*_; *V*_*ISM*_ = *V*_*ion*_ + *V*_*m*_). Thus, to obtain *V*_*ion*_, *V*_*m*_ must be measured, ideally from the same position in the cell.

The most common practical implementations of ISMs entail the use of single- or double-barrel micropipettes. For single-barrel ISMs, V_m_ is measured with a second independent (sharp) microelectrode, while for double-barrel ISMs, one channel is filled with an appropriate aqueous saline and used to measure *V*_*m*_. An obvious advantage of double-barrel ISMs is that only one impalement is required. However, their fabrication is somewhat cumbersome. Single-barrel ISMs are easier to fabricate, but more skill and instrumentation is required to impale a cell with two microelectrodes. With multicellular preparations, it is necessary to confirm that both electrodes are inside the same cell by passing bridge-corrected current pulses through the reference electrode. Configurations such as these are commonly used to measure (“normal” or “physiological”) ion activities in preparations maintained at physiological conditions, or even in live animals (Zeuthen and Monge 1975). These approaches have inherent limitations. Membrane damage due to impalement of double-barrel electrodes or two single-barrel electrodes causes membrane depolarization (Kessler, Clark et al. 1976, Koryta and Stulik 1983, Mooney, Lyall et al. 1988) which, as demonstrated below, can cause voltage-dependent re-distribution of ions of interest. In general, it is not possible to maintain a fixed membrane potential by passing current through the reference electrode because this can produce spurious voltage drops across the electrode. Any readout errors from the reference electrode will impact the quantification of ion activity. In addition, experiments to explore the effect of membrane potential on specific transport mechanisms are not possible.

To overcome these limitations, we developed a novel approach to measure dynamic changes in the intracellular activity of transported ions in intact, isolated skeletal muscle fibers under current- or voltage-clamp control. For studies of electrogenic transport processes, this method allows simultaneous measurement of ion currents and any accompanying change in intracellular ion activities; for electroneutral transport processes, it provides measurement of the intracellular ion activity changes under conditions when the membrane potential is controlled. The ability to control the membrane potential with voltage- or current-clamp eliminates contributions from any interacting voltage-dependent membrane processes.

We impale a single fiber with 4 microelectrodes. Typically, one ISM and three standard intracellular microelectrodes are used, two to measure *V*_*m*_ and one to inject current (alternatively, two different ISMs can be used). The ISM is monitored using a dual-channel high-impedance amplifier compatible with high-resistance ISMs; membrane current and voltage are controlled by a second, high-voltage two-microelectrode voltage-clamp amplifier. The ability to fix the membrane potential allows the study of factors that both depend on, and alter, membrane potential, such as occurs with electrogenic ion-transport mechanisms. In addition, the combined measurement of (net) *I*_*m*_ and myoplasmic ion activity aids identification of the charge carrier.

We demonstrate this approach using examples that address specific questions about Cl^-^, H^+^ and Na^+^ ion transport and homeostasis in skeletal muscle. The method can be easily extended to study other physiologically-relevant ions in other cell preparations.

A preliminary abstract of this work has been published (Heiny, Cannon et al. 2018).

## METHODS

### Isolated muscle fibers

Studies were performed on short (<450 μm) fibers enzymatically dissociated from flexor digitorum brevis (FDB) and interosseous muscles of C57BL mice, following previously-described protocols (DiFranco, Herrera et al. 2011, DiFranco and Vergara 2011, Fu, Struyk et al. 2011). In brief, mice were euthanized by deep anesthesia followed by cervical dislocation. An excised muscle was pinned to the bottom of a 3.5 cm Petri dish lined with Silgard and was enzymatically digested using 4mg/ml type-2 collagenase (Worthington Biochemical Corporation, USA) dissolved in HEPES-buffered Tyrode. Digestion was carried out for 35 min using a shaking water bath at 35 °C and 1 rpm. To release free fibers, the digested muscles were forced to pass back and forth through progressively smaller bore fire-polished Pasteur pipettes.

Single dissociated fibers were plated onto a 100 μm thick coverslip that formed the bottom of a 35 mm petri dish. To increase adhesion, the dissociated free fibers were washed 4-5 times using HEPES-Tyrode before plating (50-100 fibers). The fibers were kept in Tyrode solution at RT until use, and typically used within 1-8 hours after dissociation.

The experimental chamber (∼300 μl) was continuously perfused at ∼5 ml/min using a gravity-driven perfusion circuit. The solution level in the chamber was maintained constant by gentle suction. A steady solution level was found to be critical for maintaining a stable baseline current in voltage-clamp mode and for stability of potentials measured with the high-impedance amplifier for ion-sensitive microelectrodes. Solution exchanges were performed manually using multiport valves (Hamilton) in the perfusion circuit and the chamber was completely exchanged within 1 sec, as calibrated with a fluorescent dye. The standard recording solution was HEPES-buffered Tyrode, in mM: 140 NaCl, 4 KCl, 2 CaCl_2_, 1 MgCl_2_, 10 glucose, 10 HEPES, pH = 7.4 with NaOH, not gassed. For the pH studies in Figure 4, a bicarbonate-buffered Tyrode’s solution was used, which consisted of (in mM): 118 NaCl, 4.75 KCl, 2.54 CaCl_2_, 1.18 MgSO_4_, 1.18 NaH_2_PO_4_, 24.8 NaHCO_3_, 10 glucose, pH = 7.4 when gassed with 95% O_2_ and 5% CO_2_. The internal solution for the voltage-sensing microelectrodes (*V*_*m*_ and *V*_*ref*_), as well as the current-passing microelectrode (*I*_*m*_), was comprised of (in mM): 70 K-aspartate, 5 di-Na ATP, 5 di-tris creatine phosphate, 40 EGTA, 20 MOPS, 10 MgCl_2_, pH = 7.4 with KOH.

**Figure 4.**
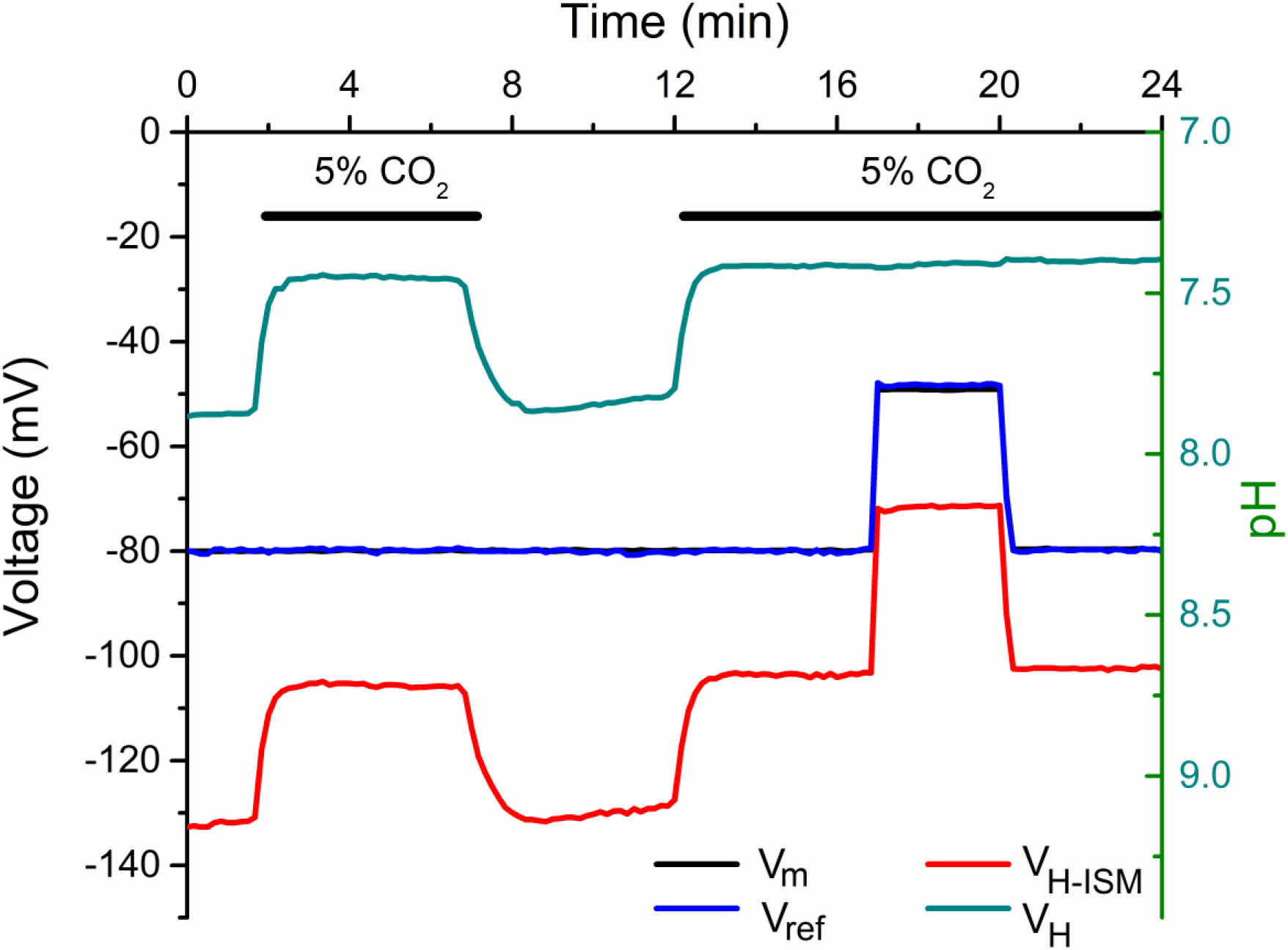
Intracellular pH rapidly responds to CO_2_ but is insensitive to a step depolarization of *V*_*m*_. Traces show simultaneous recordings of *V*_*m*_ (black), *V*_*ref*_ (blue), and *V*_*H*−*ISM*_ (red) from a fiber perfused with HEPES-buffered Tyrode’s solution (pH 7.4, nominally 0 CO_2_) that was intermittently exchanged with Tyrode’s solution containing 24 mM HCO_3_^-^ and gassed with 5% CO_2_, pH 7.4. The fiber was voltage-clamped at −80 mV and, before impalement, *V*_*H*−*ISM*_ was zeroed in the presence of Tyrode adjusted to pH = 7.0. The proton potential, *V*_*H*_ = *V*_*H*−*ISM*_ − *V*_*m*_ (green), was computed digitally off-line and is proportional to pH (right ordinate). Intracellular pH was alkaline (∼7.9) in CO_2_/HCO_3_^-^ - free Tyrode solution and rapidly acidified to 7.4 in the presence of CO_2_. A 30 mV step depolarization of *V*_*m*_ produced an identical shift of *V*_*H*−*ISM*_, therefore the difference between these two potentials remained constant (green trace, 16 to 20 min), indicating no change of *V*_*H*_ with a constant pH.

All procedures involving animals were approved by the UCLA Research Animal Committee.

### The 4-microelectrode system

The 4-microelectrode setup (Figure 1) was configured by combining a two-electrode voltage-clamp (TEVC) and a high-impedance electrometer with both a reference electrode and an ion-selective microelectrode (ISM) inputs. With this system, the intracellular activity of one or two separate ions could be measured in dissociated muscle fibers maintained under either current- or voltage-clamp conditions.

**Figure 1.**
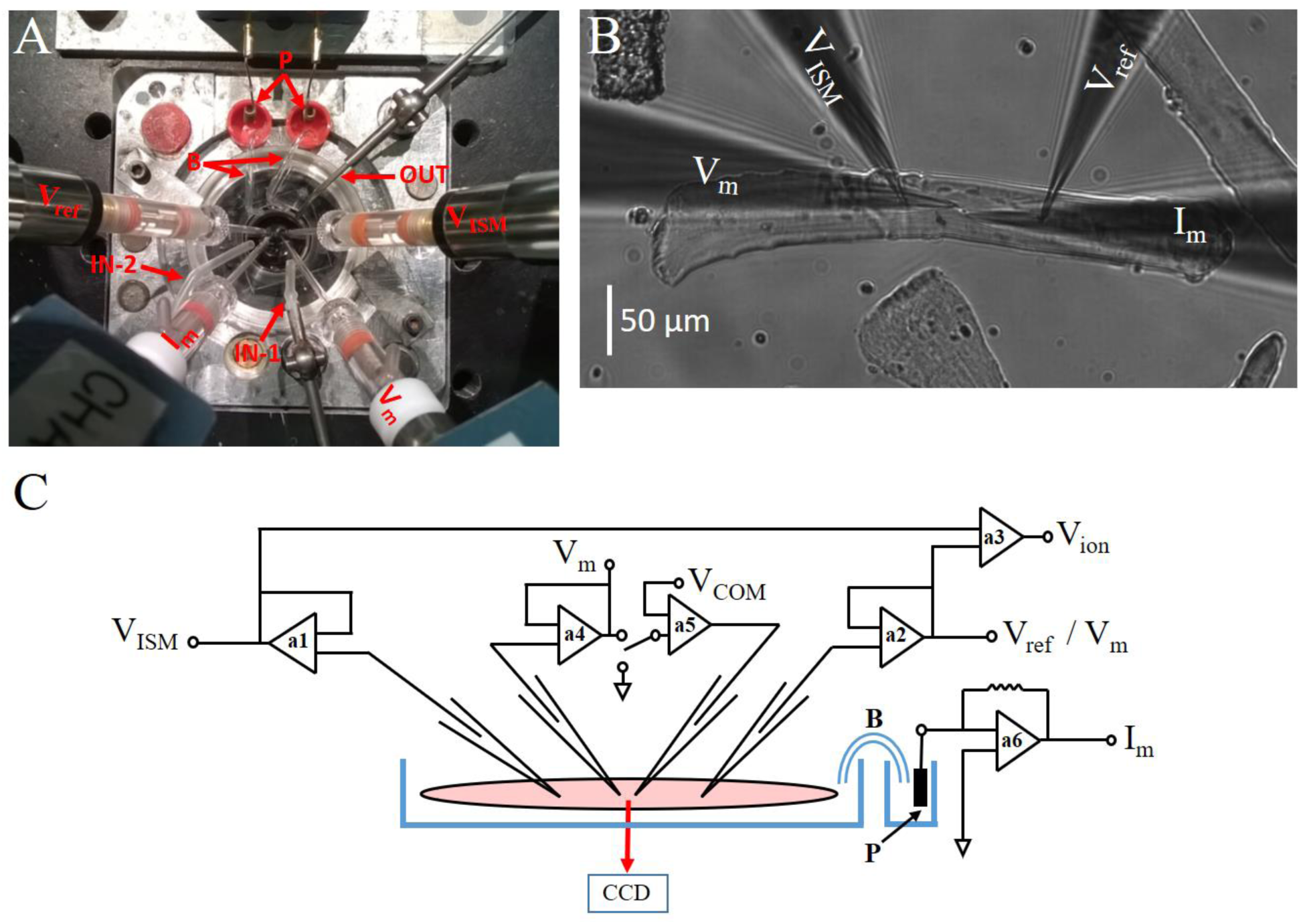
A 4-microelectrode method to measure intracellular ion activity in a voltage-clamped skeletal muscle fiber. (A) Photograph of the recording chamber on the stage of an inverted microscope. An ISM (*V*_*ISM*_) and reference microelectrode (*V*_*ref*_) were connected to a high input impedance amplifier to measure intracellular activity of Na^+^, H^+^, or Cl^-^. A second voltage-sensing microelectrode (*V*_*m*_) and current-passing microelectrode (*I*_*m*_) were used for the TEVC. The Ag/AgCl pellets (P) from the bath amplifier were connected to the recording chamber with salt bridges (B). IN-1 and IN-2 are independent solution inlets from two perfusion systems. Out is the suction outlet. (B) Photomicrograph of a typical dissociated single fiber impaled with four microelectrodes. (C) Schematic diagram of the circuit used to measure *V*_*ISM*_, *V*_*ref*_, *V*_*m*_, and *I*_*m*_. A high-voltage TEV-200A amplifier (Dagan) was used to measure and control membrane potential and membrane current. An FD223a two-channel high-impedance (10^15^Ω) amplifier (WPI, USA) was used to measure the voltage of two ion-sensing micro-electrodes (ISM) or the voltage of one ISM and one standard sharp intracellular reference microelectrdoe (*V*_*ref*_). a1, a2: high input pre-amplifiers for ISMs. a3: instrumentation amplifier for real-time calculation of *V*_*ion*_ = *V*_*ISM*_ − *V*_*ref*_. a4: membrane potential recording pre-amplifier of the voltage clamp. a5: control amplifier connected to the current injection electrode. a6: bath control amplifier (for simplicity, only on salt bridge is shown).

A TEVC with a high compliance of 145 volts (TEV-200A amplifier; Dagan, USA) was used to optimize the clamp speed of fibers up to 450 μm in length (C_m_ ∼ 2nF). Capacitance compensation was performed for both the voltage-sensing (*V*_*m*_) and current-passing (*I*_*m*_) microelectrodes of the TEVC before fiber impalement. The TEV-200A active bath clamp was used to maintain the bath at virtual ground in either current- or voltage-clamp modes. The sensing and current injection terminals of the bath amplifier were connected via salt bridges (with floating Pt wires) to the bath solution. A two-channel amplifier (FD223a; World Precision Instruments, USA) with high-impedance inputs (> 10^15^ Ω, shunted by 0.5 pF) was used to measure the voltage of two ISMs or the voltage of one ISM and one standard sharp intracellular reference microelectrode (*V*_*ref*_). No capacitance compensation was possible for *V*_*ref*_ because the device lacks this function. ISM electrodes cannot be compensated for capacitance. The ground terminals of both amplifiers were connected. For data to show fast transients in Figure 2, the voltage signals (*V*_*m*_, *V*_*ref*_, *V*_*Na*−*ISM*_) were low-pass filtered at 10 kHz, the current (*I*_*m*_) was filtered at 5 kHz, and the data were sampled at 30 μsec. Records of several minutes were used for all other Figures, and the sampling interval was 20 msec. When recording small-amplitude Na,K-ATPase currents (Figure 6), *I*_*m*_ was filtered at 100 Hz to enhance the on-line detection of current amplitude.

**Figure 2.**
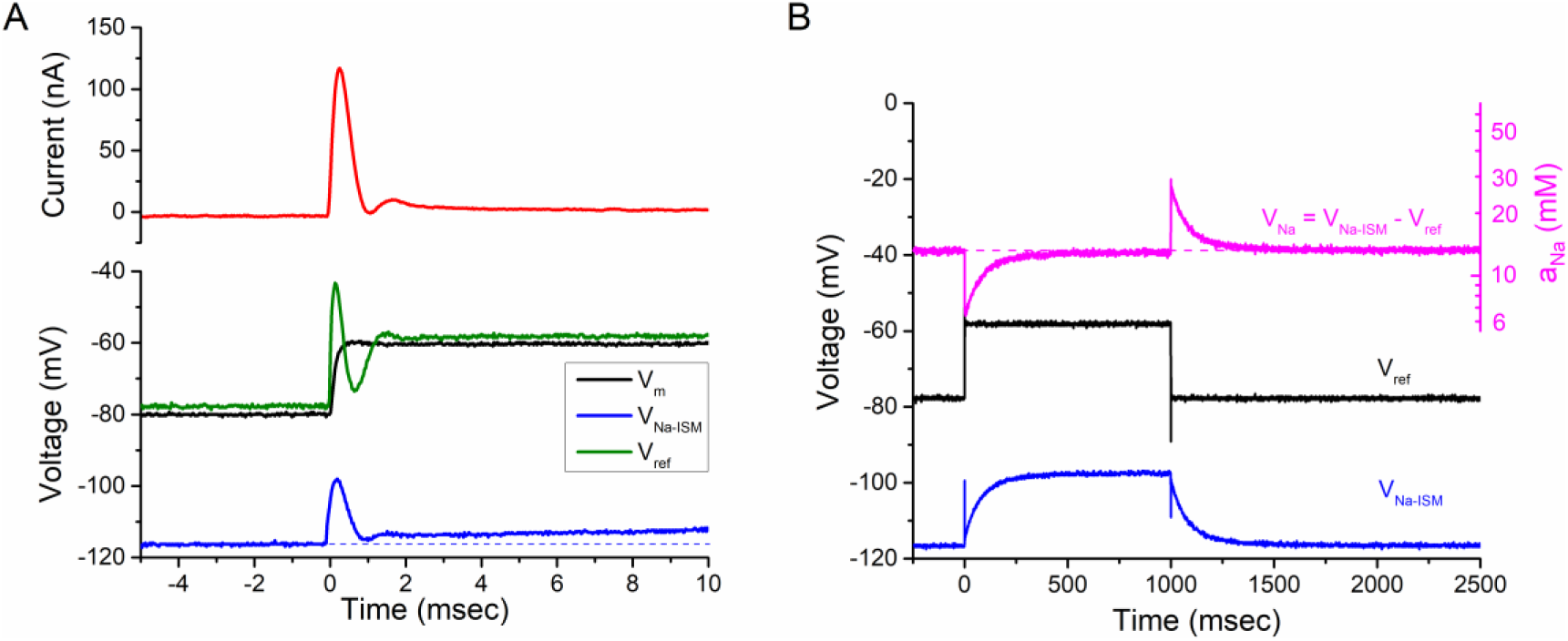
Settling time for *V*_*Na*−*ISM*_ under voltage clamp. (A) The voltage-clamp response for a 20 mV depolarization from −80 mV is shown on a fast time scale. The current transient (top panel, red) settled within 1 ms and had only a small oscillation (1 to 1.5 ms). Simultaneous recordings from the 3 voltage-sensing microelectrodes (bottom panel) shows that the rapid step change in *V*_*m*_ was competed in less than 1 ms (*V*_*m*_, black). The voltage signals from the high-impedance amplifier (*V*_*ref*_ green, *V*_*Na*−*ISM*_ blue) had larger oscillations for the first 2 ms because this device has no analog capacitance compensation. The response time of the *V*_*Na*−*ISM*_ is slow, as shown by the depolarizing shift relative to the dashed line and does not reach steady-state during this 10 ms trace. (B) A long-duration depolarization of 20 mV shows the settling time of the *V*_*Na*−*ISM*_, which in this example had a time constant of 80 ms. The intracellular *a*_*Na*_ did not change in response to the depolarization, as shown by the stable *V*_*Na*_ = *V*_*Na*−*ISM*_ − *V*_*ref*_, corresponding to an activity of 13.2 mM (dashed line, magenta).

The headstages of these amplifiers (pre-amplifiers a1, a2, a4, and a5 in Figure 1) were attached to four micromanipulators (MPC285, Sutter Instruments) that were mounted on a custom stage of an inverted fluorescence microscope (IX-100, Olympus). The angle of approach was oriented with the TEVC microelectrode pair being collinear with the fiber axis, and the two ISM microelectrodes both approaching from the same side of the fiber (Figure 1). A standard video camera was attached to the trinocular port of the microscope and used for continuous observation on a monitor during microelectrode placement and impalement at various magnifications (10x, 40x and 100x Oil). A CCD camera (Santa Barbara Instrument Group) attached to the lateral port of the microscope was used to obtain digital still images of whole impaled fibers (at 10x) for offline calculation of geometrical parameters (diameter, length, and surface area) using a custom algorithm.

### Ion-selective microelectrodes (ISM)

Micro-pipettes for both sharp and ion-sensing electrodes were pulled from 1.5 mm borosilicate capillaries with internal micro-filaments (BF150-86-10, Sutter Instruments). A 4-step pulling protocol using a horizontal puller (P97, Sutter Instruments) was optimized to obtain a short shaft (∼8 mm) leading to a sub-micron tip opening. The electrodes had a tip resistance of 10-12 MΩ when filled with internal solution (typically 100 mM monovalent salts). The use of capillaries with micro-filaments significantly speeded the manufacture of ISMs without compromising the sealing of the organic phase to the inner face of the capillary wall. Capillaries were used as sold, without additional cleaning.

Silanized micro-pipettes must be used for ISM fabrication because the hydrophobic surface is essential for mechanical stability and high electrical resistance at the interface between glass and the liquid membrane at the tip of the electrode. Micropipettes were silanized by evaporation of chlorotrimethylsilane (Sigma) in a 500 ml jar, followed by baking at 250 °C for at least 4 h, as previously described (Difranco, Quinonez et al. 2019). Silanized micro-pipettes were stored dry in a closed jar (WPI Jar-E215) to avoid dust adhesion and were used for up to 2 weeks without noticeable deterioration.

Ion-selective microelectrodes were assembled by backfilling through a 34G microneedle, as previously described (Difranco, Quinonez et al. 2019). Briefly, an excess of the ionophore-containing cocktail was delivered as closely as possible to the micro-pipette tip. Within a few minutes, the mixture spontaneously wicks into the tip and the excess is removed with a new clean microneedle until a layer only 200 to 300 μm remains. The saline solution is then backfilled with another microneedle, taking care to insure a clean interface with the organic layer and no bubbles. Prior to use, each ISM was individually tested using the calibration solutions. Acceptable electrodes were stored in sealed jars (WPI Jar-E215) with tips submersed in filtered (0.2 μm) saline. ISMs were used up to 2 weeks after fabrication, with no detectable deterioration.

The ionophore cocktails were mixed and stored as stock solutions at room temperature that could be used for up to 1 year. The composition (w/w %) of the ionophore and the backfilling saline solutions were:

Sodium microelectrode (Na-ISM): 10% Na^+^ ionophore IV (Sigma-Aldrich, #71745), 1% Na-tetraphenylborate in NPOE. Backfilled with 100 NaCl, 10 HEPES, pH = 7.4 titrated with HCl

Chloride microelectrode (Cl-ISM): 10% mercuracarborand-3 (MC3, custom synthetized (Difranco, Quinonez et al. 2019)), 2.5% tridodecylmethyl ammonium chloride (TDMAC), in 2-nitrophenyl octyl ether (NPOE). Backfilled with 100 NaCl, 10 HEPES, pH = 7.4 titrated with H_2_SO_4_.

Proton (pH) microelectrode H-ISM: 10% H^+^ ionophore-I (Sigma-Aldrich #95292), 0.7% tetraphenylborate in NPOE. Backfilled with 100 NaCl, 100 HEPES, pH = 7.0 with HCl.

### Calibration of Ion-selective Microelectrodes

A three-point calibration check was used immediately after ISM fabrication to select high-quality electrodes. We verified that all three types of ISMs had linear Nernstian responses over the physiological ranges of interest (1 to 100 mM for Na^+^ and Cl^-^, 6.0 to 8.0 pH units for H), wherein *V*_*ion*−*ISM*_ was proportional to the logarithm of ion activity, *log*(*a*_*ion*_).

The calibration solutions, prepared as molar concentration, were converted to activity for the ion of interest, *a*_*ion*_, by using the extended Debye-Hückel equation (Davies 1962), which accounts for both ionic strength and ionic radius. For the Na-ISM, calibration solutions of 1, 10, and 100 mM Na^+^ were prepared by mixing 150 Na^+^ stock solution (in mM: 150 NaCl, 7.2 TEA-OH, 10 HEPES, pH = 7.4 with TEA-OH) and 160 NMDG stock solution (in mM: 160 NMDG, 10 HEPES, pH = 7.4 titrated with HCl). For the Cl-ISM, calibration solutions of 1, 10, and 100 mM Cl^-^ were prepared by mixing 150 Cl^-^ stock solution (in mM: 140 NaCl, 4 KCl, 2 CaCl2, 1 MgCl_2_, 10 HEPES, pH = 7.4 with NaOH) and 0 Cl^-^ stock solution (in mM 115 Na_2_CO_4_, 2 K_2_SO_4_, 2 Ca(OH)_2_, 1 Mg(OH)_2_, 10 HEPES, pH = 7.4 titrated with H_2_SO_4_. For the H-ISM, calibration solutions of varying pH were prepared from a Tyrode’s stock solution for which the buffer at 10 mM was either MES (pH = 6.0 or 6.5), HEPES (pH = 7.0 or 7.5), or Tris (pH 8.0).

The offset of each ISM was set to 0 mV in the following solutions to facilitate the on-line interpretation of ionic activity during an experiment: 100 mM Na-Tyrode for Na-ISM, 100 Cl-Tyrode for Cl-ISM, HEPES buffered pH 7.0 Tyrode for H-ISM.

For intracellular measurements, the voltage of the ion-selective microelectrode is *V*_*ion*−*ISM*_ = *V*_*ion*_ + *V*_*m*_, where *V*_*ion*_ is the electrochemical potential detected by the selective ionophore and *V*_*m*_ is the membrane potential of the cell. The value of *V*_*m*_ used for calculating *a*_*ion*_ was measured as the potential detected by the reference microelectrode (*V*_*ref*_) of the high-impedance amplifier. Digital subtraction was used to compute *V*_*ion*_ = *V*_*ion*−*ISM*_ − *V*_*ref*_. Finally, ion activity was calculated as 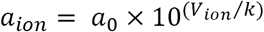, where *a*_0_ is the activity of the control solution when *V*_*ion*_ was set to 0 mV and *k* is the slope of the linear relation between *V*_*ion*_ and *log*(*a*_*ion*_). At the end of the experiment, the ISM was withdrawn from the fiber and stability of the three-point calibration was verified with the same solutions used for the initial screen of ISM quality.

## RESULTS

### Fiber impalement and quality of the voltage-clamp

We have previously demonstrated that high-quality voltage-clamp recordings can be achieved in intact, short (< 450 μm) mammalian skeletal muscle fibers by using a high-voltage two-microelectrode voltage clamp (DiFranco, Herrera et al. 2011, DiFranco and Vergara 2011, Fu, Struyk et al. 2011). This approach can be used to obtain reliable measurements of the major ionic currents (DiFranco, Herrera et al. 2011, DiFranco and Vergara 2011, Fu, Struyk et al. 2011, DiFranco, Quinonez et al. 2012, DiFranco, Yu et al. 2015) and transport mechanisms (DiFranco, Hakimjavadi et al. 2015) in skeletal fibers. Here, we introduce an approach that simultaneously adds measurement of intracellular ion activity with measurement of transmembrane ion fluxes under voltage-clamp conditions. To achieve this, we extended our method for voltage clamping short fibers by adding a two-channel high-impedance amplifier compatible with ion-sensing microelectrodes.

A top view of the 4-electrode configuration is shown in Figure 1A, demonstrating the placement of microelectrodes, salt bridges, and perfusion components around the experimental chamber mounted on the stage of the inverted microscope. Ideally, the microelectrodes would be oriented orthogonally from each other to reduce stray electrical interactions. The actual orientation that we achieved is smaller (∼60°) due to space constraints from the two salt bridges and virtual-ground pre-amplifier. The salt bridges connect the bath solution to the voltage measuring and current output of the virtual ground amplifier, and are essential to limit changes in junction potentials upon bath solution changes.

The *V*_*m*_ and *I*_*m*_ microelectrodes of the TEVC were located near the center of the fiber with a separation of about 20 μm and the ion-sensing microelectrode pair, *V*_*ISM*_ and *V*_*ref*_, were placed more laterally at approximately 100 μm from each end of the fiber (Figure 1B). Clean electrode impalements with minimal damage to the fiber were achieved with the following protocol. First, the TEVC microelectrodes were inserted into the fiber. Small current pulses (4-5 nA, 5 ms) were applied at 3 Hz through the *I*_*m*_ electrode in current-clamp mode while it was slowly advanced towards the fiber. Contact with the membrane was signaled by an increase in the voltage drop at the electrode tip (i.e. an increase in tip resistance). A small additional advance of the microelectrode led to membrane puncture, as evident from a sudden change in steady potential (between −45 to −65 mV) and a reduction of the voltage drop during the pulse. Fiber impalement with the *V*_*m*_ microelectrode was performed next, and again an abrupt change in voltage plus the membrane charging in response to the injected current pulse were used to detect membrane penetration rather than a change in visual appearance. A small holding current (<10 nA in high-quality fibers of length up to 450 μm) was then applied in current-clamp mode to bring the resting potential to −80 mV. As the fiber recovered over the next 3 to 5 min, the holding current required to maintain a *V*_*m*_ of −80 mV usually decreased.

Next, the ion-sensing microelectrodes were inserted into the fiber. Impalement with the *V*_*ref*_ microelectrode was detected by an abrupt voltage change from 0 mV to within 2 or 3 mV of the holding potential of −80 mV, as reported by the *V*_*m*_ electrode. Confirmation was obtained by measuring the *V*_*ref*_ voltage transient in response to a 5 nA current pulse applied through the *I*_*m*_ microelectrode. Finally, an ISM (for Na^+^, Cl^-^, or H^+^) was inserted into the fiber. Impalement was detected as a voltage jump, recognizing that the amplitude is the sum of *V*_*ion*_ + *V*_*m*_, which was small in the case of a Cl^-^ electrode (e.g. −80 + 65 = −15 mV) and was very large for the Na^+^ or H^+^ electrode (e.g. −80 + (−40) = −120 mV for Na^+^, as shown in Figure 2).

Several criteria were used to define a successful 4-electrode preparation. First, the initial holding current should be small (< 10 nA) to maintain a holding potential of −80 mV. Second, the holding current should not increase more than 50% with the subsequent insertion of the ISM microelectrodes. Third, *V*_*m*_ and *V*_*ref*_ should agree within 3 mV. Fourth, *V*_*ISM*_ should be near the expected value of *V*_*ion*_ + *V*_*ref*_. Finally upon withdrawal from the fiber at the end of the experiment, the voltage-sensing microelectrodes (*V*_*m*_ and *V*_*ref*_) should read 0 ± 3 mV, and a repeat of the 3-point calibration of the *V*_*ISM*_ should be within ± 3 mV of the values obtained before impalement.

### Response time of an ion-selective microelectrode in a voltage-clamped fiber

One goal of the present work was to demonstrate that an electrically-stable condition can be achieved when short FDB fibers, already impaled with two sharp microelectrodes connected to the voltage clamp amplifier, are also impaled with 1 or 2 additional ISMs connected to another (high-impedance) amplifier. Conversely, we verified that the ISM performance is not adversely affected by the voltage-clamp amplifier working in either current- or voltage-clamp configuration.

The response of an FDB fiber impaled with four microelectrodes (*V*_*m*_ and *I*_*m*_ for TEVC, plus *V*_*ref*_ and *V*_*Na*−*ISM*_) is shown in Figure 2. The fiber was held at −80 mV in voltage-clamp mode. The holding current was small (−3.5 nA) and did not drift, which shows that the fiber was healthy and without signs of damage from microelectrode impalement. As expected, the *V*_*Na*−*ISM*_ signal (blue trace) was more negative than *V*_*m*_ because the ISM was initially zeroed in a 100 mM Na^+^ solution, whereas internal *a*_*Na*_ is about 10 mM. The quality of the voltage clamp is shown by the response to a 20 mV step depolarization on a fast time scale (Figure 2A). The clamp speed was comparable to the performance we see in fibers studied with a conventional TEVC alone (DiFranco, Herrera et al. 2011, Fu, Struyk et al. 2011). With a clamp feedback gain of ∼75%, both the capacitance transient of *I*_*m*_ (red trace) was complete and *V*_*m*_ (black trace) reached steady-state in less than 1 ms and the measured membrane potential change (*V*_*m*_, 10 kHz bandwidth) closely followed the imposed 20 mV step. The voltage clamp was stable, without large oscillations detected in *V*_*m*_ or *I*_*m*_. The potentials measured with the FD223a amplifier had transient biphasic oscillations (*V*_*Na*−*ISM*_ blue trace, *V*_*ref*_ green trace) at the onset of the pulse, most likely because these electrodes cannot be compensated for capacitance. Importantly, the voltage detected by *V*_*ref*_, placed ∼100 μm from *I*_*m*_, was comparable to *V*_*m*_ sensed by the voltage clamp and attests to the quality of the space clamp. Therefore, we are confident that the local membrane potential at the site of the ion-sensing microelectrode is also equal to *V*_*m*_. This constancy is essential because *V*_*ion*_ is derived from *V*_*ion*−*ISM*_ − *V*_*ref*_, and the corresponding interpretation of ion activity is exponentially related to this difference. In practice, *V*_*ref*_ was always within a few mV of *V*_*m*_, which implies the *V*_*ref*_ microelectrode can be omitted to reduce the complexity of the system (and potential damage to the fiber). Alternatively, a second *V*_*ion*−*ISM*_ can be used so that two different intracellular ion activities can be measured simultaneously or the second *V*_*ion*−*ISM*_ can be placed just outside the fiber to obtain a direct measure of the local transmembrane ion gradient. A slowly-depolarizing drift was observed for *V*_*Na*−*ISM*_ after the pulse onset (Figure 2A, blue trace) because, on the short time scale of 10 ms, the response had not reached steady-state.

The response to a long-duration voltage step is shown for a different FDB fiber in Figure 2B. The response of the Na-ISM (blue trace) exponentially approached steady-state with a time constant of about 80 ms. This lag in the *V*_*Na*−*ISM*_ response is typical for a high-impedance ISM, as contrasted with the fast response of the reference microelectrode *V*_*ref*_ (black trace) to measure membrane potential. Because the settling times of these two microelectrodes are different, the subtraction used to compute *V*_*Na*_ = *V*_*Na*−*ISM*_ − *V*_*ref*_ will transiently be invalid after each voltage jump. This effect is shown by the *V*_*Na*_ signal (magenta trace) in Figure 2B. At the onset of the 20 mV depolarization (t = 0 ms), the difference signal, *V*_*Na*_, had an apparent decrease that returned to baseline within 500 ms (i.e. 4 time constants of *V*_*Na*−*ISM*_), as would be expected for a constant intracellular Na^+^ activity. The mirror-image effect occurs at the off edge of the pulse. While this temporal bandwidth limitation of ISMs is well-known (Ammann 1986), our 4-microelectrode setup provides a new method to directly measure the ISM settling time while the electrode is in the intracellular environment. Alternatively, the voltage-clamp can be used to hold *V*_*m*_ constant, even in the face of bath solution exchanges, drug application, or other perturbations, which ensures that a change in the *V*_*ion*−*ISM*_ signal is caused by a change of intracellular *a*_*ion*_ and not a shift of the membrane potential.

### Voltage-dependent changes of intracellular Cl^-^ activity

The membrane properties of skeletal muscle enabled us to test the ability of our 4-microelectrode setup to detect a voltage-dependent shift of intracellular ion activity. The chloride conductance is especially large in resting muscle, about 1 mS/cm^2^ (Dulhunty 1979, Pedersen, de Paoli et al. 2009), whereas the active transport of Cl^-^ by cotransporters and exchangers is small in comparison (∼15 pmol/cm^2^) (Geukes Foppen 2004). Because of this arrangement, the equilibrium potential for Cl^-^, *E*_*Cl*_, is only a few mV depolarized from the membrane potential (Aickin, Betz et al. 1989). If a sustained change of *V*_*m*_ is imposed, then the prediction is that intracellular Cl^-^ will shift until *E*_*Cl*_ is again only a few mV depolarized relative to *V*_*m*_. While indirect evidence in support of this concept has been provided (Aickin, Betz et al. 1989), there never has been a direct measurement showing a change of intracellular *a*_*Cl*_ in response to an imposed shift of *V*_*m*_.

The intracellular *a*_*Cl*_, measured from an FDB fiber with our 4-ME setup in voltage-clamp mode, is shown in Figure 3. As before, the *V*_*m*_ response of the clamp and the *V*_*ref*_ signal were nearly identical (black and blue traces, respectively, in top panel). The voltage sensed by the Cl-ISM was close to 0 mV (red trace) because *E*_*Cl*_ was only a few mV depolarized from *V*_*m*_. On the time scale of hundreds of seconds in this figure, the slow transients in the *V*_*Cl*−*ISM*_ trace were caused by shifts of intracellular *a*_*Cl*_ (not the settling time of the ISM which is less than 1 second, Figure 2). Also note that for this anion-selective ISM, the voltage becomes more negative as the intracellular *a*_*Cl*_ increases. The change of intracellular *a*_*Cl*_ is shown more clearly by computing *V*_*Cl*_ = −(*V*_*Cl*−*ISM*_ − *V*_*ref*_), as shown in the bottom panel of Figure 3 (green trace). Finally, at the end of the record (700 to 1100 sec), the Cl-ISM was withdrawn from the fiber and the electrode calibration was repeated. These data clearly show that substantial shifts of intracellular *a*_*Cl*_ occur in intact muscle fibers when a sustained shift of *V*_*m*_ is imposed. Moreover, these data provide the first measure for the time course of the *a*_*Cl*_ shift, which had a time constant on the order of 3 to 10 s.

**Figure 3.**
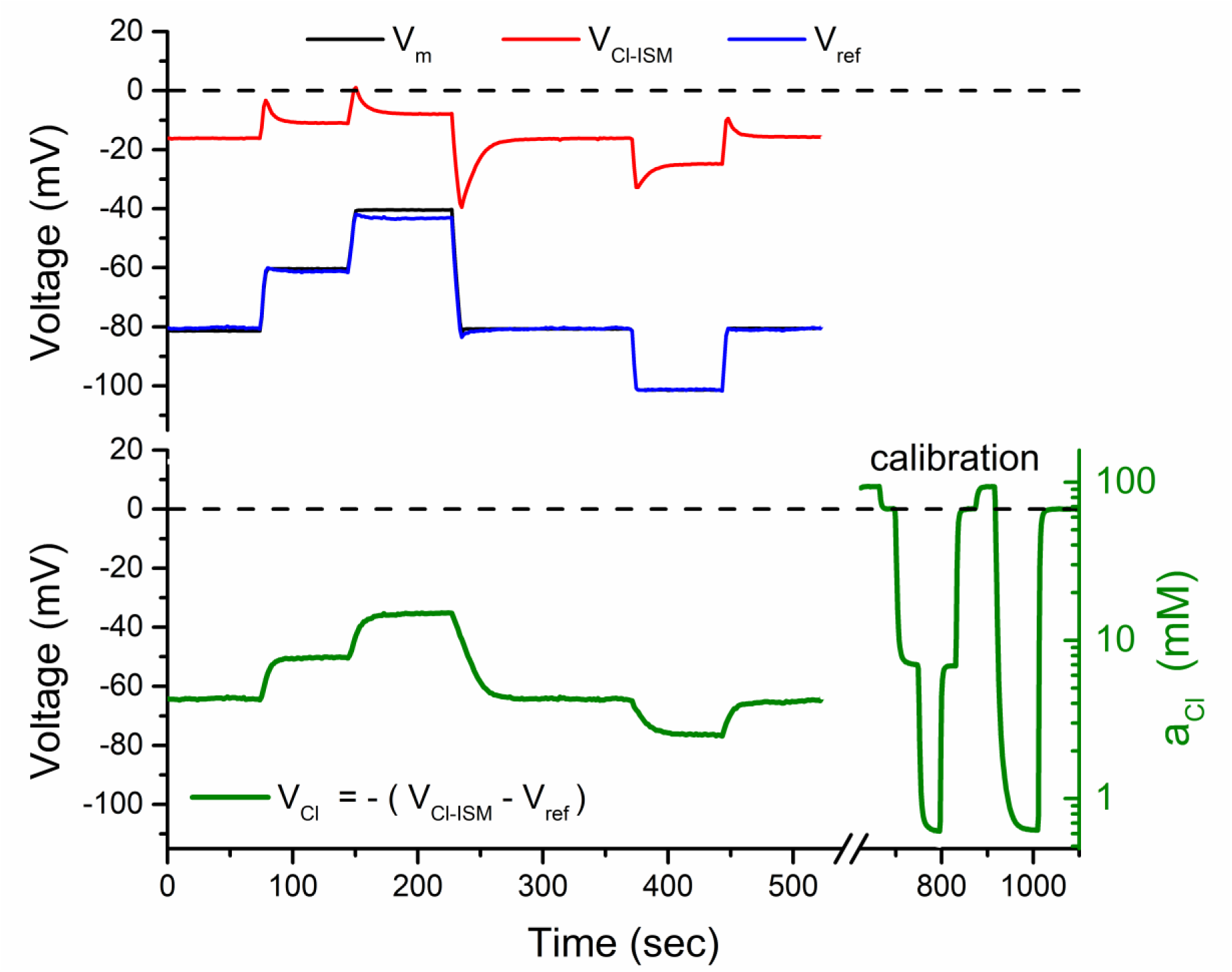
Voltage-dependent movement of chloride ions. Upper panel shows simultaneous recordings of *V*_*m*_ (black), *V*_*ref*_ (blue), and *V*_*Cl*−*ISM*_ (red) from a fiber under voltage clamp. The Cl-ISM was initially zeroed in modified Tyrode’s solution with a [Cl^-^] of 100 mM, corresponding to *a*_*Cl*_ = 69 mM. The Cl potential, *V*_*Cl*_ = −(*V*_*Cl*−*ISM*_ − *V*_*ref*_), was computed off-line and is plotted in the lower panel, along with the calibration response for the Cl-ISM that was performed after the electrode was withdrawn from the fiber. Fiber depolarization induced a substantial increase of intracellular *a*_*Cl*_ with a time constant of ∼5 s and was fully reversible, albeit slower (time constant ∼10 s), with repolarization of the fiber. The chamber was continuously perfused with HEPES-buffered Tyrode’s solution.

### A pH-selective microelectrode detects pCO_2_ changes and is insensitive to membrane potential

We used a pH-sensitive ISM with the 4-microelectrode setup to demonstrate a shift of intracellular *a*_*H*_ produced by exchange of the bathing solution and also to show that the difference signal, *V*_*H*_ = *V*_*H*−*ISM*_ − *V*_*ref*_, was insensitive to an imposed voltage jump of *V*_*m*_. The fiber was initially held at −80 mV in voltage-clamp mode, as shown in Figure 4. Again, the *V*_*m*_ and *V*_*ref*_ signals were indistinguishable (black and blue traces, respectively) and the *V*_*H*−*ISM*_ potential (red trace) was more negative than *V*_*hold*_ because this ISM was zeroed in a pH 7.0 calibration solution. The difference signal, *V*_*H*_ = *V*_*H*−*ISM*_ − *V*_*ref*_ (green trace), is proportional to intracellular pH, as shown by the ordinate scale on the right in Figure 4. The control bath solution was HEPES-buffered Tyrode (pH = 7.4) that contained no bicarbonate and was not gassed with CO_2_, which results in intracellular alkalosis as shown by the initial pH of about 7.85 (interval 0 to 2 min). Bath exchange with a 24 mM HCO_3_-buffered Tyrode’s solution that was pre-equilibrated with 5% CO_2_ / 95% O_2_ (pH 7.4) caused a rapid decrease of intracellular pH to 7.45 (Figure 4, interval 2 to 7 min) that was reversible upon return to HCO_3_/CO_2_-free Tyrode’s solution.

A second bath exchange back to 5% CO_2_ was performed at 12 min, followed by a 30 mV step depolarization (Figure 4, 17 to 20 min). The *V*_*H*−*ISM*_ signal had a nearly-identical step change in voltage (red trace, 17 to 20 min), on this time scale of minutes for which the settling time for the ISM was much faster. Consequently, the difference signal which yields *V*_*H*_ had no significant change (green trace), as expected when the intracellular pH remains constant.

### Demonstration of electrically silent and electrogenic Na^+^ transport in a voltage-clamped fiber

The combined ability to measure ion activity with an ISM and to voltage-clamp the cell is a powerful approach that can be used to study a variety of transport mechanisms. We show an example where the 4-microelectrode setup was used to measure intracellular Na^+^ accumulation/depletion in response to modulation of Na,K-ATPase pump activity imposed by changing external [K^+^]. Our prior voltage-clamp studies showed that in isolated fibers, *I*_*pump*_ can be stimulated by increasing external [K^+^] from 4 to 20 mM (20K condition) or stalled by removing K^+^ from the external solution (0K condition), and that this process is reversible (DiFranco, Hakimjavadi et al. 2015). It is predicted that these changes in pump activity will cause commensurate changes in intracellular Na^+^, which in turn will modulate pump activity; however, this has not been directly measured in skeletal muscle.

Figure 6 shows, for the first time, simultaneous measurements of intracellular *a*_*Na*_ and *I*_*pump*_ in a fiber voltage-clamped at −85 mV while pump activity was modulated by step changes of external [K^+^] from 0 mM to 20 mM. When the pump was stalled in the 0K condition, intracellular Na^+^ rapidly increased, as shown by the depolarized shift of *V*_*Na*−*ISM*_ (Fig. 6A red trace). The Na-ISM potential was converted to *a*_*Na*_ and plotted in Figure 6B to illustrate the substantial magnitude of this intracellular Na^+^ accumulation. This Na^+^ load was then rapidly cleared when the Na,K-ATPase was re-activated by a return to the 20K condition. The process was fully reversible and could be repeated over several cycles, which demonstrates the stability of the preparation even with a fiber impaled by 4 microelectrodes.

As we previously reported from studies with a traditional TEVC (DiFranco, Hakimjavadi et al. 2015), a large transient pump current was rapidly activated by the transition from 0K to 20K (Fig. 6C). This net outward current reflects the 3 Na^+^ : 2 K^+^ stoichiometry of the Na,K-ATPase. The decay of *I*_*pump*_ correlates with the reduction of intracellular *a*_*Na*_, demonstrating the well-known regulatory effect of Na_i_ on pump activity. In contrast to the clear electrogenicity of Na^+^ clearance by the Na,K-ATPase, Na^+^ loading while the pump was stalled in 0K was almost silent electrically. Only a brief inward current surge was observed at the onset of the 0K condition. Presumably most of the net Na^+^ entry occurs via an electroneutral exchanger or cotransporter (but not the NKCC1 cotransporter, which would be inactive in 0K, nor Na/HCO_3_ exchangers in this bicarbonate-free preparation).

These data also provide a new method to characterize the dependence of pump activity on intracellular *a*_*Na*_. The data from Figures 6B and 6C are redrawn as a phase-plot of *I*_*pump*_ vs *a*_*Na*_ in Figure 6D. Starting from the lower left corner of the plot (low *a*_*Na*_ in 20K), the transition to 0K causes Na^+^ loading with very little change in *I*_*m*_ (horizontal trajectory to the right). From here, the transition back to 20K produces an abrupt increase in *I*_*pump*_ (steep upward segment). The high Na,K-ATPase activity then causes intracellular *a*_*Na*_ to decrease (leftward movement), which in turn decreases *I*_*pump*_ (downward movement) until the trajectory returns to the starting point. This upper limb of the closed-loop, labeled Na^+^ clearance in the plot provides a direct measure of Na,K-ATPase activity as a function of intracellular *a*_*Na*_ for a given constant external [K^+^].

## DISCUSSION

### Combined voltage-clamp and ISM recording is robust in muscle fibers

Our experience with a new 4-microelectrode technique firmly establishes the feasibility of simultaneously using an ISM to measure intracellular ion activity while also performing voltage-clamp recordings from dissociated muscle fibers. The addition of the *V*_*ISM*_ and *V*_*ref*_ electrodes did not compromise the clamp speed nor cause the clamp to become unstable and oscillate (Figure 2). Furthermore, the close agreement of *V*_*ref*_ (ion-sensing circuit) and *V*_*m*_ (TEVC circuit) confirmed the high quality of the space-clamp in these short fibers. Viewed from the alternative perspective, we showed that the quality of the ISM recording was not adversely impacted by the voltage clamp. The extraordinarily high impedance of the ISM circuit makes the system susceptible to noise from electromagnetic activity in the environment. In our experience, however, the output of the ISM circuit, *V*_*ion*_ = *V*_*ion*−*ISM*_ − *V*_*ref*_, remained stable and free from artifact while the fiber was voltage-clamped and even when a step change of *V*_*m*_ was imposed (Figure 4).

The fiber impalement with 4 microelectrodes was well tolerated and the muscle preparation remained viable with stable responses often lasting an hour or longer. Several criteria support our interpretation that fibers remained healthy in the 4-microelectrode configuration. First, the resting potential remained relatively constant (± 3 mV), even as the second and third microelectrodes were inserted (*V*_*rest*_ cannot be measured in isolation for the fourth, ISM, electrode). Second, in voltage-clamp mode, only a small current (∼ −5 nA) was required to hold *V*_*m*_ at −80 mV and this current did not drift over time. The preparation was stable enough to reliably measure small Na,K-ATPase pump currents of 5 to 10 μA/cm^2^ over a 30 min interval (Figure 5). Third, the measured activity of intracellular Na^+^, H^+^, and Cl^-^ remained constant over tens of minutes, under conditions where *V*_*m*_, pump activity and bath composition were held constant, implying that there were no substantial leakage currents that would dissipate ion gradients.

**Figure 5.**
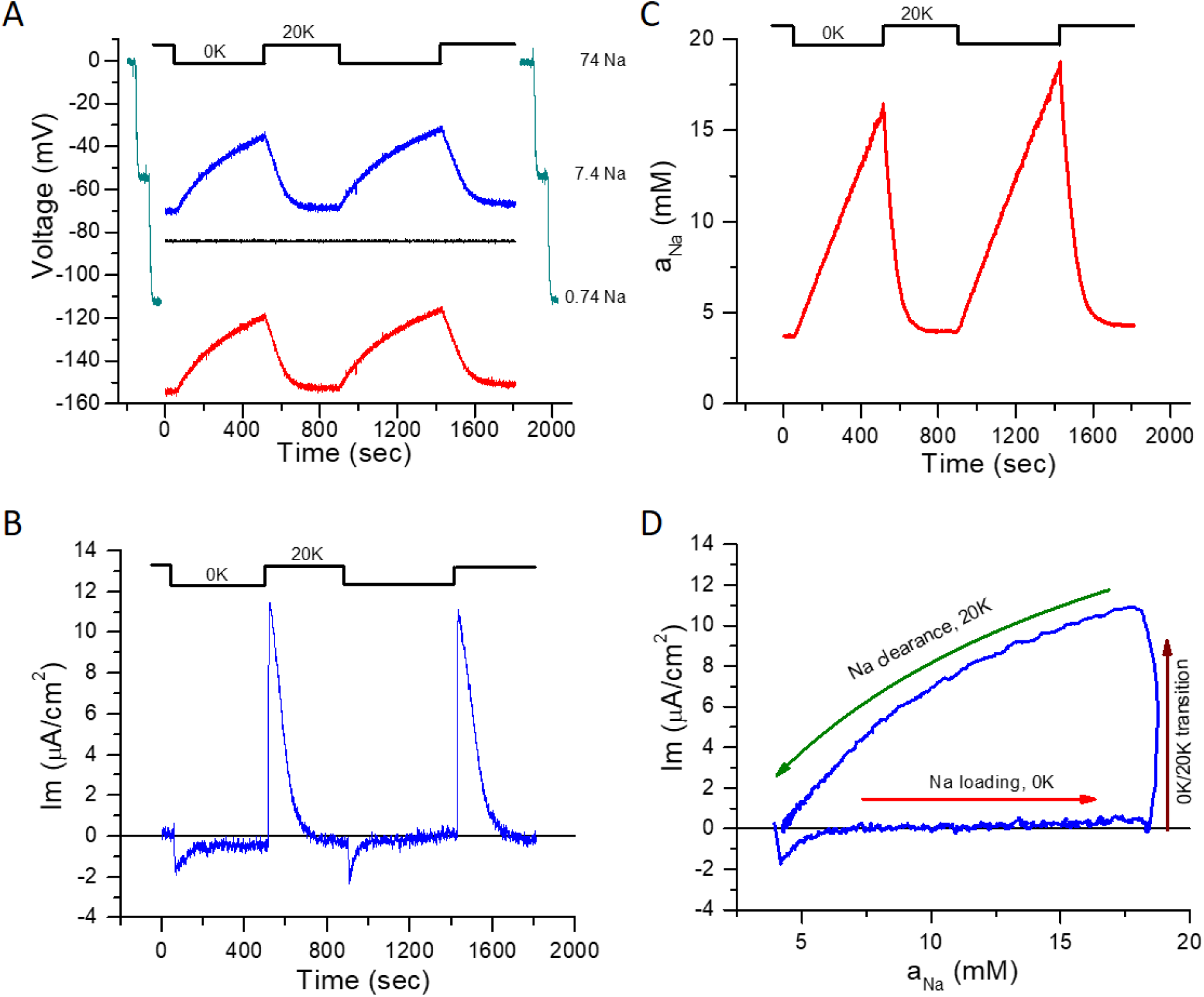
Passive and active sodium translocation at a constant membrane potential. (A) Simultaneous records of *V*_*m*_ (black trace) and *V*_*Na*−*ISM*_ (red trace) from a voltage-clamped fiber maintained at a holding potential of −85 mV. Extracellular [K^+^] was switched between 0 and 20 mM (compensated by replacement with Na^+^) in a continuously-perfused chamber. The Na-dependent potential (blue trace) was computed as *V*_*Na*_ = *V*_*Na*−*ISM*_ − *V*_*m*_. Inhibition of the Na,K-ATPase in 0 mM K^+^ caused a rise of [Na]_i_ that was revered in 20 mM K^+^. *V*_*ref*_ was indistinguishable from *V*_*m*_ and is omitted. The traces shown at each side of the intracellular records represent the pre-(left) and post-calibration (right) traces of the Na-ISM. (B) Membrane current density recorded simultaneously with the voltage records in (A). Electrogenic extrusion of intracellular Na^+^ by the Na,K-ATPase produced a large outward current transient. *I*_*m*_ was zeroed in the presence of 20 mM K^+^. K^+^, Ca^2+^ and Cl^-^ currents were blocked using Ba^2+^ (1 mM), nifedipine (20 μM) and 9ACA (200 μM), respectively. (C) Intracellular Na^+^ activity calculated from *V*_*Na*_ = *V*_*Na*−*ISM*_ − *V*_*m*_. (D) Phase plot of membrane current density as a function of the intracellular Na^+^ activity. Data are the responses in the time interval 800 to 1800 s in (B) and (C).

Prior studies with self-referencing ISMs constructed from double-barreled theta glass capillaries often showed signs of fiber damage with low resting potentials and unstable recordings with drift (Aickin and Brading 1982, Kondo, Igarashi et al. 1993). Our design with separate *V*_*ref*_ and *V*_*ISM*_ electrodes allows for smaller tip sizes than typically used with theta glass, which may account for the improved fiber viability. Our multiple (single-barrel) electrode technique for measuring intracellular ion activity while also in voltage-clamp could be adapted for use in other preparations with large cells such as *Xenopus* oocytes (Virkki, Wilson et al. 2002) or invertebrate neurons.

### Technical advances from using the 4-microelectrode system

The ability to measure intracellular ion activity with ISMs in a voltage-clamped fiber is an important advance for the study of ion gradients in skeletal muscle and provides new opportunities to investigate ion transport systems. Of the available methods to measure intracellular ion content, the ISM technique has several advantages: (i) a large linear operating range, approximately 4 orders of magnitude; (2) the most relevant property, ion activity, is directly measured; (3) measurement of absolute activity, not relative; (4) stability over tens of minutes, very little drift; (5) highly-selective ionophores are available for most ions of interest; (6) low cost and equipment requirements. Another advantage is that the ISM method provides a simultaneous measure of *V*_*m*_, and now in the 4-electrode configuration *I*_*m*_ is measured as well.

On the other hand, fiber impalement with microelectrodes may cause a run-down of *V*_*rest*_. If the ion gradient is sensitive to *V*_*m*_, as we show here for Cl^-^ (Figure 3), then the depolarization from impalement may cause an unwanted change of intracellular ion activity. The addition of the voltage clamp avoids this concern and adds new capabilities to the recording system. First, the voltage clamp can be used to maintain *V*_*m*_ at the value of *V*_*rest*_ determined from the initial impalement of the fiber. The ability to maintain a constant *V*_*m*_ is also advantageous when studying the effects of perturbations (e.g. drugs or changes in external ion concentration) that would otherwise change *V*_*rest*_ and thereby confound the interpretation of a change in intracellular ion activity. Alternatively, *V*_*m*_ can be set to any desired value or voltage steps can be applied to study voltage-dependent responses of intracellular ion activity. As shown in Figure 2, this technique can be used to experimentally verify the settling time of the ISM and thereby define the high-frequency cut-off for reliably measuring *V*_*ion*_ = *V*_*ion*−*ISM*_ − *V*_*ref*_ when *V*_*m*_ is not constant.

Another advantage of the voltage-clamp configuration is the measurement of total membrane current, *I*_*m*_, while simultaneously measuring intracellular ion activity. For example, the value of *I*_*m*_ needed to maintain any desired *V*_*hold*_ can be used as a criterion to ensure fiber viability after impalement of all the microelectrodes. Consequently, *I*_*m*_ also provides information about net ion flux which can be combined with ISM measurement of ion activity to characterize membrane transport systems. Here, we show that the combination of measuring dynamic changes of intracellular Na^+^ activity and *I*_*pump*_ can be used to characterize the intracellular Na-dependent of the Na,K-ATPase in an intact skeletal muscle fiber (Figure 5).

## Abbreviations

9ACA: 9-anthracene carboxylic acid;
BMT: bumetanide;
ISM: liquid ion-selective microelectrode;
MC3: [9]-mercuracarborand-3;
NMDG: N-methyl-D-glucamine;
NPOE: 2-nitrophenyl octyl ether;
TDMAC: tridodecylmethyl ammonium chloride;
TEVC: two-electrode voltage-clamp.

## ACKNOWLEDGEMENT

This work was supported by grants R01-AR063710 (NIAMS / NIH to JH), R01-AR63182 (NIAMS / NIH to SC), and R21-AR067422 (NIAMS / NIH to MD) and by the Muscular Dystrophy Association (MDA RG 381149 to SC). We thank Dr. J. Lopez Padrino for kindly sharing his knowhow on the fabrication of liquid membrane ion-selective microelectrodes.

